# ACSS2 contributes to transcriptional regulation in Cajal-Retzius cells in a mouse model of Alzheimer’s disease

**DOI:** 10.1101/2024.04.08.588604

**Authors:** Gabor Egervari, Desi C. Alexander, Greg Donahue, Hua Huang, Connor Hogan, Mariel Mendoza, Benjamin A. Garcia, Nancy M. Bonini, Shelley L. Berger

**Author notes:** These two authors contributed equally to this work (Gabor Egervari, Desi C. Alexander).

## Abstract

Dysregulation of histone acetylation in the brain has emerged as a major contributor to human Alzheimer’s disease (AD). The mechanisms by which these protective or risk-conferring epigenetic marks are established and maintained are under intense investigation. ACSS2 (Acetyl-CoA Synthetase 2) is a key metabolic enzyme that is chromatin-associated in neurons. ACSS2 is recruited to specific promoters and generates a local pool of acetyl-CoA from acetate, thereby fueling histone acetylation and driving the expression of neuronal genes that regulate learning and memory. Here, we examine the contribution of ACSS2-mediated histone acetylation to AD-related molecular and behavioral outcomes. Using a mouse model of human pathological AD-Tau injection, we show that loss of ACSS2 exacerbates Tau-related memory impairments, while dietary supplementation of acetate rescues learning in an ACSS2-dependent manner. Combining state-of-the-art proteomic and genomic approaches, we demonstrate that this effect is accompanied by ACSS2-dependent incorporation of acetate into hippocampal histone acetylation, which facilitates gene expression programs related to learning. Further, we identify Cajal-Retzius neurons as a critical hippocampal neuronal population affected, exhibiting the largest epigenetic and transcriptional dysregulation. Overall, these results reveal ACSS2 as a key neuroprotective metabolic enzyme, dysregulation of which might play an important role in the etiology of human AD, and guide the development of future therapies for AD and related dementia.

## INTRODUCTION

Recent evidence for an intricate interaction between metabolic and epigenetic processes has led to a fundamental shift in models of transcriptional regulation. In particular, nuclear metabolic enzymes have emerged as central players in epigenetic regulation^1,2^. Contrary to prevailing dogma, these enzymes are directly recruited into the nucleus and to chromatin, where they generate a local, transient supply of metabolites that serve as substrates or cofactors for histone and DNA modification enzymes. This metabolic-epigenetic interface is rapidly emerging as a potential area for novel therapeutic interventions in a variety of conditions linked to epigenetic and transcriptional dysregulation.

We and others have reported marked changes of histone acetylation in the brain of Alzheimer’s disease (AD) patients^3–6^. For example, while normal aging is characterized by increased acetylation of H4K16, AD brains reveal a striking loss of this modification^3^. This pattern suggests that H4K16 acetylation has a protective effect during normal aging that is lost in AD brains. On the other hand, acetylation of H3K27 and H3K9 are enriched in the brains of AD patients, particularly at genes related to chromatin and disease pathways, implicating these acetylation sites as potentially risk conferring for AD^4^.

Nevertheless, general elevation of histone acetylation has been shown to improve memory across multiple mouse AD models^7–10^. Administration of broad-spectrum HDAC inhibitors (HDACi) elevates histone acetylation levels in the hippocampus and cortex, and improves associative and spatial memories^7^. In addition to behavioral impacts, HDACi administration increases dendritic spine number^7^, reduces phosphorylated Tau levels^10^, and restores long term potentiation of synaptic activity^8^. These results suggest that broadly increasing histone acetylation levels could confer neuronal resilience against disease.

The molecular mechanisms that govern the AD-specific reconfiguration of the histone acetylation landscape of the hippocampus may be central to disease. Here, we assess the contribution of ACSS2 (Acetyl-CoA Synthetase 2), a novel metabolic regulator of brain histone acetylation^11–13^ to epigenetic changes in tau-associated AD models. In neurons, ACSS2 is predominantly nuclear^11^. Moreover, ACSS2 is required for histone acetylation and expression of specific neuronal genes, allowing for hippocampal learning and memory^11,12,14^. AD is characterized by both amyloid plaques and tau neurofibrillary tangles, but tau-associated dysfunction is more directly linked to disease progression^15–17^ Several models for tau toxicity and pathology have been generated in the mouse by injection of human AD-associated Tau aggregates into the mouse brain^18^. These models show pathology that strikingly mimics the human disease, making such models exceptional for capturing pathological aspects of human AD in mice.

Here we show that AD-Tau seeding recapitulates key molecular changes of human AD, and define ACSS2-mediated histone acetylation as a novel modulator of AD-Tau related molecular and behavioral phenotypes: loss of ACSS2 exacerbates memory impairments, while dietary supplementation of acetate, the substrate of this enzyme, rescues learning and tau-related transcriptional deficits. Combining state-of-the-art proteomic and genomic approaches, we show that this is accompanied by ACSS2-dependent incorporation of acetate into hippocampal histone acetylation to facilitate gene expression programs that drive learning. Using single nuclei approaches, we identify Cajal-Retzius neurons as a critical hippocampal neuronal population affected. This work outlines new, therapeutically targetable mechanisms underlying the offset and progression of AD that are independent of traditional efforts focusing on Tau accumulation and amyloid plaque formation.

## METHODS

### Mice

Animal use and all experiments performed were approved by the Institutional Animal Care and Use Committee (IACUC, protocols 804849). All personnel involved have been adequately trained and are qualified according to the Animal Welfare Act (AWA) and the Public Health Service (PHS) policy. To prevent genetic drift, Acss2^KO^ and corresponding WT controls were generated through homozygous crosses of F1 progeny from heterozygote breeding cages. Mice were housed on a 12/12h light/dark cycle (7 am to 7 pm), with food and water provided ad libitum. To investigate the effects of chronic acetate treatment, mice were maintained on LabDiet 5001 with 5% Na-acetate or calorically equivalent control (manufactured and irradiated by TestDiet, Land O’Lakes Inc.). All behavioral experiments were conducted between 7 am and 11 am to reduce time-of-day effects.

### Purification of pathological tau protein from human AD brains

Tau protein was purified as previously described^19^. Briefly, 6–14 g of frontal cortical gray matter from AD patients was homogenized using a Dounce homogenizer in nine volumes (v/w) of high-salt buffer (10 mM Tris-HCl, pH 7.4, 0.8 M NaCl, 1 mM EDTA, and 2 mM dithiothreitol [DTT], with protease inhibitor cocktail, phosphatase inhibitor, and PMSF) with 0.1% sarkosyl and 10% sucrose added and centrifuged at 10,000 g for 10 min at 4°C. Pellets were reextracted once or twice using the same buffer conditions as the starting materials, and the supernatants from all two to three initial extractions were filtered and pooled. Additional sarkosyl was added to the pooled low-speed supernatant to reach 1%. After 1h nutation at room temperature, samples were centrifuged again at 300,000 g for 60 min at 4°C. The resulting sarkosyl-insoluble pellets, which contain pathological Tau, were washed once in PBS and then resuspended in PBS (∼100 μl/g gray matter) by passing through 27-G 0.5-in needles. The resuspended pellets were further purified by a brief sonication (20 pulses at ∼0.5 s/pulse) using a hand-held probe (QSonica) followed by centrifugation at 100,000 g for 30 min at 4°C, whereby the majority of protein contaminants were partitioned into the supernatant, with 60–70% of tau remaining in the pellet fraction. The pellets were resuspended in PBS at one fifth to one half of the precentrifugation volume, sonicated with 20–60 short pulses (∼0.5 s/pulse), and spun at 10,000 g for 30 min at 4°C to remove large debris. The final supernatants were used in the study and referred to as AD Tau. The use of postmortem brain tissues for research was approved by the University of Pennsylvania’s Institutional Review Board with informed consent from patients or their families.

### AD-Tau injections

6 months old male and female WT and ACSS2 KO mice were anaesthetized with isoflurane gas (1–5% to maintain surgical plane) and placed in a sterile field within a stereotaxic device. Artificial tears were applied to eyes to ensure sufficient lubrication. Animals received an injection of bupivacaine (2.5 mg kg−1) for local anaesthesia before the skin was disinfected with betadine solution and the skull exposed with a short incision using sterile surgical equipment. A small hole (about 0.5 mm) was drilled in the skull over the target area using a stereotax and a stereotactic drill. A micro-syringe filled with AD Tau lysate was inserted into the right dorsal hippocampus and slowly removed following injection of 1 μg Tau (AP, − 2.5 mm; DV, − 1.4 mm; ML, + 2 mm from bregma). All animals received a single dose of subcutaneous meloxicam (5 mg/kg) as analgesia at induction and one dose per day for two days postoperatively as needed. To allow sufficient time for the development of AD-like pathology, mice underwent behavioral or molecular characterization 6 months post-injection.

### Fear conditioning

Mice were handled for three consecutive days for 2 minutes each prior to the start of the experiment. On the day of fear acquisition, mice were individually placed in conditioning chambers (Med Associates) and habituated to the novel environment for 2.5 minutes prior to delivery of a 2s long, 1.5 mA foot shock. Mice were promptly removed 30s after shock onset and returned to their home cages. Chambers were wiped down with 70% ethanol between each round. 24 hours later, freezing response was tested for 5 consecutive minutes using FreezeScan^TM^ software (CleverSys, Inc).

### Object location memory

Non-aversive spatial memory was assessed as previously described^12,20^. Briefly, mice were handled for 2 minutes a day for three consecutive days (KO). On training day, mice were habituated to an empty arena for 6 minutes, then exposed to a collection of three distinct objects (glass bottle, metal tower, plastic cylinder) of roughly similar sizes for 6 minutes. This training was repeated three times, with mice removed and all surfaces wiped down with 70% ethanol between trainings. For testing, the mice were returned to the same arena 24 hours later, this time with one object moved to a new location. Mice were allowed to explore freely for 6 minutes. Mice were monitored using a video camera, and time spent interacting with each object was assessed afterwards using Any-maze (Stoelting Co.). Quantification of mouse-object interactions was blinded. Discrimination index was calculated according to [DI = (t_displaced_ – [(t_stationary1_ + t_stationary2_)/2])/(t_displaced_ + [(t_stationary1_ + tstationary2)/2) × 100]]

### Open field

Mice were placed in an open arena (30 cm x 40 cm) and allowed to explore freely for 6 minutes. Resulting videos were analyzed using ANY-maze behavioural tracking software (Stoelting Co.). Locomotion was assayed using path length (in meters) over the entire 6 minute period. Thigmotaxis (a measure of anxiety) was measured by quantifying the amount of time spent within the peripheral zone (defined as 8 cm from the arena walls) over the entire period.

### Elevated zero maze

The maze was a circle-shaped platform of five centimeters in diameterwith two opposing open arcs and two opposing closed arcs of equal size, elevated 30 cm above the ground^21^.. Mice were placed in the middle of a closed arc and were allowed to freely explore the maze for 5 minutes. Each test was videotaped and the behavior of the rat was subsequently scored by an independent and blinded observer. Arm entry was defined as entering an arm with all four paws. At the end of the 5 min test period, the mice were removed from the maze, the floor was wiped with a damp cloth, and any faecal boluses removed. The percentage of time spent in the open arcs [time spent in open arcs/(time spent in open arcs + time spent in closed arcs)×100] and the percentage of the number of open-arc entries [open arcs entries/(open arcs entries +closed arcs entries)×100] were used as a measure of anxiety.

### Y maze

The Y-maze test of spontaneous alternation assess working spatial memory in rodents^22^. Mice were gently placed in the distal portion of an arm in a Y-shaped arena and allowed to explore freely. Mice were recorded by video cameras, and arm entries were scored by eye by blinded investigators. Arm entries were called when the mouse’s hindquarters passed fully into the new arm. Percent spontaneous alternations were quantified according to the formula [SAR = [Number of alternations/(total arm entries – 2)] x 100.

### Histology and immunohistochemistry

Animals were transcardially perfused with 1× PBS followed by 4% paraformaldehyde in PBS. Following dissection, brain tissues were transferred to PBS and subsequently embedded in paraffin and sectioned at a thickness of 5 μm. Every tenth section was stained with hematoxylin and eosin (H&E), and sections from similar anatomical planes were chosen for histologic and immunohistochemical analyses. Following deparaffinization and rehydration, sections were stained with phosphor-Tau (Ser202, Thr205) monoclonal antibody clone AT8 (Thermo, cat# MN1020) to detect Tau phosphorylation. Histopathologic analysis was performed in a blinded manner.

### Histone extraction, propionylation and digestion

Histones were extracted from 1 mm^2^ punches taken from the dorsal CA1 region of the mouse hippocampus by using Nuclei Isolation Buffer (NIB) as previously described^23,24^. The tissue punches were homogenized using biovortexer in NIB (15 mM Tris-HCl, 15 mM NaCl, 60 mM KCl, 5 mM MgCl_2_, 1 mM CaCl_2_ and 250 mM sucrose at pH 7.5; 0.5 mM AEBSF, 10 mM sodium butyrate, 5 µM microcystin and 1 mM DTT added fresh) with 0.3% NP-40 on ice, and washed twice in NIB without NP-40 to remove the detergent from the nuclear pellet. Next, pellets were incubated in 0.2LM H_2_SO_4_ for 4 hours, and the supernatant was collected after centrifugation for 5 min at 3400 x g. Finally, histones were precipitated with 25% trichloroacetic acid (TCA) overnight. The histone pellet was then washed with ice-cold acetone-0.1% HCl and acetone twice to remove residual TCA. Histones were derivatized and digested as previously described (30). Histone pellets were resuspended in 30 µl of 50 mM ammonium bicarbonate (pH 8.0), and 15 μl derivatization mix was added to the samples, which consist of propionic anhydride and acetonitrile in a 1:3 ratio (v/v), and this was immediately followed by the addition of 7.5 µl ammonium hydroxide to maintain pH 8.0. The sample was incubated for 15 min at 37 °C, dried and the derivatization procedure was repeated one more time to ensure complete derivatization of unmodified and monomethylated lysine residues. Samples were then resuspended in 50 µl of 50 mM ammonium bicarbonate and incubated with trypsin (enzyme:sample ratio of 1:20) overnight at room temperature. After digestion, the derivatization reaction was performed again twice to derivatize the N-termini of the peptides. Samples were desalted using C_18_ stage tips before LC–MS analysis and dried. Finally, the peptides were resuspended in 0.1% formic acid (FA) prior to nLC-MS/MS.

### Mass spectrometry

Samples were analyzed by using a nanoLC-MS/MS setup. NanoLC was configured with a 75 μm ID x 25 cm Reprosil-Pur C18-AQ (3 μm; Dr. Maisch GmbH, Germany) nano-column using an EASY-nLC nano-HPLC (Thermo Scientific, San Jose, CA, USA), packed in-house. The HPLC gradient was as follows: 5% to 32% solvent B (A = 0.1% formic acid; B = 80% acetonitrile, 0.1% formic acid) over 45 minutes, from 32% to 90% solvent B in 5 minutes, 90% B for 10 minutes at a flow-rate of 300 nL/min. nanoLC was coupled to an Orbitrap Elite mass spectrometer (Thermo Scientific, San Jose, CA, USA). The acquisition method was data independent acquisition (DIA) as described. Briefly, two full scan MS spectra (m/z 300−1100) were acquired in the ion trap within a DIA duty cycle, and 16 ms/ms were performed with an isolation window of 50 Da.

Normalized collision energy (CE) was set to 35%. Raw MS data were analyzed manually. We selected the 7 most intense peptides of histone H3 and H4 containing acetylations, and we extracted the relative abundance of the M+1, M+2 and M+3 isotopes compared to the monoisotopic signal. The other peptides were not considered as, due to their low abundance, we could not reliably quantify the relative abundance of all the isotopes. The percentage represented in the radar plots indicates the relative intensity of the M+3 signal (the fourth isotope) as compared to the monoisotopic signal. Data were not normalized to the non-labeled sample, so that the relative abundance of the natural isotopic distribution can be appreciated also in the untreated mice.

### RNA extraction and RNA-sequencing

RNA was extracted and libraries were prepared as previously described^11^.. Briefly, total RNA was extracted using Trizol-chloroform from 1 mm-thick slices of dHPC (comprising the CA1, CA2, CA3, and dentate gyrus). Total RNA quality was assessed on the Bioanalyzer platform using the RNA 6000 Nano assay (Agilent). mRNA was isolated from 300 ng total RNA using the NEBNext® Poly(A) mRNA Magnetic Isolation Module (E7490L), and libraries were prepared using the NEBNext® Ultra™ II RNA Library Prep Kit for Illumina® (E7770). All RNA-seq data were prepared for analysis as follows: NextSeq sequencing data was demultiplexed using native applications on BaseSpace. Demultiplexed FASTQs were aligned by RNA-STAR 2.5.2 to assembly mm10 (GRCm38) (parameters: --outFilterType BySJout --outFilterMultimapNmax 20 --alignSJoverhangMin 8 -- alignSJDBoverhangMin 1 --outFilterMismatchNmax 999 --alignIntronMin 20 --alignIntronMax 1000000 -- alignMatesGapMax 1000000). Aligned reads were mapped to genomic features using HTSeq 0.9.1 (parameters: -r pos -s rev -t exon -i gene_id). Quantification, library size adjustment, and differential gene expression analysis were performed using DESeq2. The significance of gene alterations were determined using the Wald test with multiple test correction according to the Benjamini-Hochberg method with FDR < 0.05. Gene Ontology (GO) analysis was performed using the DAVID bioinformatics suite^25^, and top terms associated with biological processes were reported.

### ChIP-seq

ChIP-seq was performed as previously described. Briefly, ∼20 mg dorsal hippocampus from each mouse was minced on ice and crosslinked with 1% formaldehyde for 10 min and quenched with 125 mM glycine for 5 min. Nuclei were prepared by dounce homogenization of crosslinked tissue in nuclei isolation buffer (50 mM Tris-HCl at pH 7.5, 25 mM KCl, 5 mM MgCl2, 0.25 M sucrose) with freshly added protease inhibitors and sodium butyrate. Nuclei were lysed in nuclei lysis buffer (10 mM Tris-HCl at pH 8.0, 100 mM NaCl, 1 mM EDTA, 0.5 mM EGTA, 0.1% Na-deoxycholate, 0.5% N-lauroylsarcosine) with freshly added protease inhibitors and sodium butyrate and chromatin was sheared using a Covaris S220 sonicator to ∼250 bp size. Equal aliquots of sonicated chromatin were used per immunoprecipitation reaction with 4 ul H3K27ac antibody (Abcam, cat #4729; lot # GR323132-1) preconjugated to Protein G Dynabeads (Life Technologies). 10% of the chromatin was saved as Input. ChIP reactions were incubated overnight at 4°C with rotation and washed three times in wash buffer. Immunoprecipitated DNA was eluted from the beads, reversed crosslinked and purified together with Input DNA. 10 ng DNA (either ChIP or Input) was used to construct sequencing libraries using the NEBNext Ultra II DNA library prep kit for Illumina (New England Biolabs, NEB). Libraries were multiplexed using NEBNext Multiplex Oligos for Illumina (dual index primers) and single-ended sequenced (75 bp) on the NextSeq 500 platform (Illumina) in accordance with the manufacturer’s protocol. ChIP-seq tags generated with the NextSeq 550 platform were demultiplexed with the bcl2fastq utility and aligned to the mouse reference genome (assembly GRCm38/mm10) using Bowtie v1.1.1 allowing up to two mismatches per sequencing tag (parameters -m 1 --best). Peaks were detected using MACS2 (tag size = 75 bp; FDR < 1x10^-3^) from pooled H3K27ac tags of mice from the same condition along with treatment-matched Input tags as control. The MTL method^26^ was used to compare H3K27ac enrichment in the four study conditions. Statistical significance of differential H3K27ac enrichments was assessed by using DiffBind (Bioconductor v3.7) in a 2-way comparison (wt-saline vs wt-EtOH or ACSS2KD-saline vs ACSS2KD-EtOH) across the individual replicate samples (FDR < 10%). UCSC Genome Browser track views were created for ChIP-seq data by first pooling replicates and generating coverage maps using BEDtools genomeCoverageBed -bg, then adjusting for library size using the RPM coefficient. Input signal was then subtracted from chIP signal. Resulting tracks were converted from bedGraph to bigWig using the Genome Browser’s bedGraphToBigWig utility. RNA-seq tracks were created similarly, first splitting by tag orientation to the genomic reference strand and then creating coverage maps. Because an RPM adjustment might disguise a large deformation in the transcriptome distribution, maps were adjusted for library size using the average scalar coefficient size factor determined by DESeq2. Resulting tracks were converted to bigWigs as ChIP-seq tracks were, and the + and - tags from a given sample were plotted as overlays in a track hub.

### Single nuclei RNA-seq and ATAC-seq

Single-nuclei dissociation was performed following the 10X Genomics user guide for nuclei isolation from complex tissues (CG000375 Rev B). Briefly, bilateral hippocampal tissue was homogenized in NP40 Lysis Buffer (10 mM Tris-HCl pH 7.4, 10 mM NaCl, 3 mM MgCl2, 0.1% NP40, 1 mM DTT) using biovortexer on ice. Following 5 min incubation, lysates were passed through 70 um Flowmi strainer and centrifuged at 4C, 500g for 5 min. Pellets were washed and resuspended in ice-cold PBS containing 1% BSA. Nuclei were stained with ready-made 7AAD solution (Sigma, SML1633) and sorted on MoFlo Astrios EQ (Beckman Coulter). Sorted nuclei were pelleted, resuspended in 0.1x Lysis Buffer (10 mM Tris-HCl pH 7.4, 10 mM NaCl, 3 mM MgCl2, 1% BSA, 1 mM DTT, 0.01% Tween-20, 0.01% NP40, 0.001% digitonin) and incubated on ice for 2 min. Permeabilized nuclei were washed with Wash Buffer (10 mM Tris-HCl pH 7.4, 10 mM NaCl, 3 mM MgCl2, 1% BSA, 0.1% Tween-20, 1 mM DTT) and resuspended in 1x Diluted Nuclei Buffer with 1 mM DTT. All buffers contained 1 U/ul Sigma Protector RNase inhibitor (3335402001). Nuclei concentration was assessed via trypan blue staining and counting with a hemocytometer. This approach yielded approximately ∼200-300k nuclei per brain. Library preparation was performed using Chromium Next GEM Single Cell Multiome ATAC+Gene Expression kit (10x Genomics) following the user guide (CG000338 Rev C). Targeted nuclei recovery rate was 10,000 nuclei/sample. For ATAC-seq, DNA and libraries were amplified for 7 cycles using indexes from Sample Index Plate N, Set A. Library size distribution and concentration were measured using BioAnalyzer High Sensitivity DNA kit (Agilent) and NEB quantification kit. 2.7 pM of pooled libraries were loaded onto 150-cycle high-yield Illumina flowcells and sequenced on Illumina NextSeq 550 using custom recipe for read configuration 50:8:16:49. For RNA-seq, cDNA was amplified for 9 cycles and libraries were amplified for 14 cycles using indexes from Dual Index Plate TT Set A. Library size distribution and concentration were measured using BioAnalyzer DNA1000 kit (Agilent) and NEB quantification kit. 2.7 pM of pooled libraries were loaded onto 150-cycle high-yield Illumina flowcells and sequenced on Illumina NextSeq 550 using read configuration 28:10:10:90.

## RESULTS

### Injection of human pathological AD-Tau into mouse hippocampus recapitulates human AD-like molecular dysfunction

Injection of human pathological AD-Tau (Tau) isolated from post-mortem brains of Alzheimer’s disease (AD) patients induces AD-like pathology in mice characterized by hyperphosphorylation of endogenous Tau and spreading of the pathology^19,27^. Given the central role of Tau in AD, we used this model to test whether AD-associated molecular changes are recapitulated in the mouse by injecting AD-Tau into the hippocampus of wild-type C57Bl6/J mice . Tau phosphorylation occurred in the dentate gyrus of Tau-injected mice, but not in PBS-injected littermates (**Figure 1A**). We then extended this work to gene expression and histone acetylation implicated in human AD^3,4^. Brain tissue exhibited peak AD-like pathology at 6 months post-injection^19^, so we focused on this time point. We noted substantial dysregulation of H3K27ac peaks, with the majority reduced upon Tau injection (15,481 PBS-specific, 8,856 Tau-specific) (**Figure 1B,C).** While transcriptional changes were limited overall, we observed differential expression of several genes linked to AD and neurodegeneration. These included *Hpcal1* (Hippocalcin Like 1) and *Itga4* (Integrin Subunit Alpha 4), genes with known variants associated with increased AD risk^28,29^. We also observed downregulation of *Crmp1* (Collapsin Response Mediator Protein 1) and *Madd* (MAP Kinase Activating Death Domain), changes that have been linked to neuronal death^30^ and correlate with neuron loss in AD^31^. We also noted altered expression of enzymes that regulate histone acetylation, such as *Anp32a* (Acidic Nuclear Phosphoprotein 32 Family Member A), which has been linked to synapse and memory loss in mouse models of AD^32^ and *Hdac2* (Histone Deacetylase 2), which has been implicated in neurodegeneration and cognitive dysfunction^33^.

**Figure 1.**
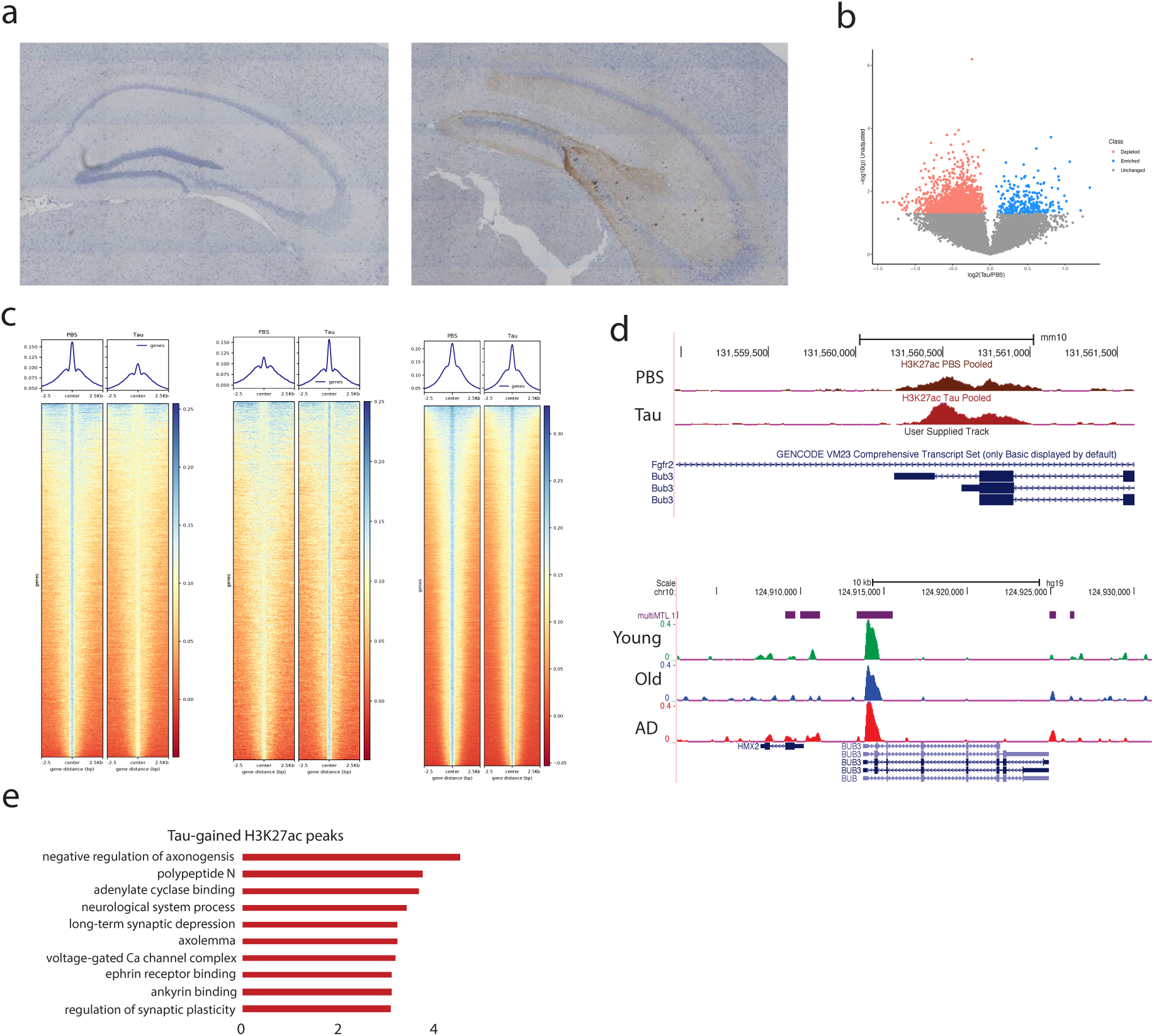
AD-Tau injection in mouse hippocampus recapitulates human AD pathology and epigenetic changes. **A.** AT8 immunohistochemistry showing endogenous Tau phosphorylation in the hippocampus of AD-Tau injected (1 μg) but not PBS-injected mice 6 months post seeding. **B.** A volcano plot showing t test -log10(p-values) vs log2(Tau/PBS) for a filtered group of H3K27ac peaks significantly different between Tau-injected and PBS control. Peaks filtered such that H3K27ac-input > 0.1 across all replicates in the biased direction. **C**. Heatmaps showing 5kb windows around all H3K27ac peaks in Tau or PBS, separated by Tau-specific, PBS-specific, or Common (N = 15,481 PBS-specific, 8,856 Tau-specific, and 27,540 common peaks). **D.**Overlapping H3K27ac enrichment at Bub3 gene in the hippocampus of AD-Tau injected mice vs PBS controls (top) and in AD patients compared to young and old controls (bottom). Gene Ontology analysis of H3K27ac peaks induced by AD-Tau injection in mouse hippocampus.

Analysis of histone H3 lysine 27 acetylation (H3K27ac) in the mouse hippocampus showed that Tau injection recapitulated changes observed in human AD, with increased acetylation at loci analogous to that of the human brain. For example, both mouse and human hippocampi showed increased H3K27ac at the transcription start site of *Bub3* (**Figure 1D**), a gene encoding for the Bub3 mitotic checkpoint protein implicated in neurodegeneration and other aging-associated phenotypes^4,34^. Genome-wide, H3K27ac peaks specific to Tau injection (top 5%) were strongly enriched at genes we previously identified as showing H3K27ac dysregulation in human AD^4^ . Gene Ontology analysis of genes associated with Tau-specific H3K27ac peaks revealed enrichment of genes related to the regulation of axonogenesis, neurological system processes, long-term synaptic depression and the regulation of axonal function and synaptic plasticity (**Figure 1E**); these processes may contribute to the AD-like learning and memory impairments in this model (see **Figure 2**).

**Figure 2.**
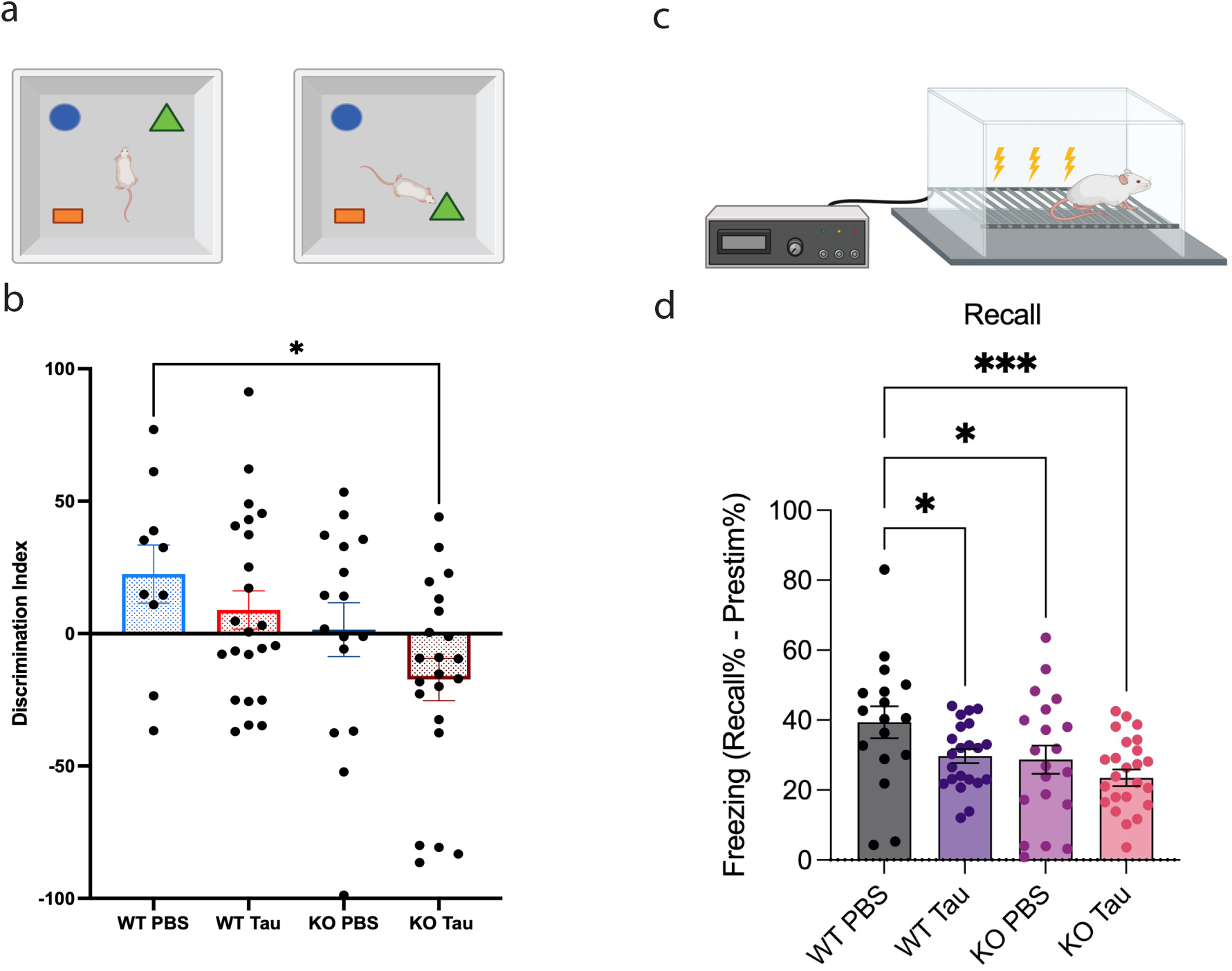
ACSS2 deletion exacerbates learning and memory impairments induced by hippocampal AD-Tau injection. **A.** Long-term memory was tested using object location memory (OLM). **B.** Decreased discrimination index of moved objects in AD-Tau injected ACSS2 KO mice indicate more severe impairments of learning compared to other groups. **C.** Contextual fear conditioning was used as an alternative assay of long-term spatial memory formation. **D.** Freezing levels showing more severe memory impairments in AD-Tau injected ACSS2 KO mice compared to other groups.

Taken together, these findings underscore AD-Tau injection as a translational mouse model for AD that recapitulates not only pathological, but also transcriptional and epigenetic changes reflective of the brain of human AD patients.

### ACSS2 knock out exacerbates learning and memory impairments induced by AD-Tau

To focus on the role of ACSS2 in AD-associated cognitive dysfunction, we performed AD-tau injections in WT and ACSS2 KO mice and performed a battery of behavioral tests 6 months post-injection to characterize learning and memory impairments associated with hippocampal Tau injection. As reported previously for Tau injection^27^, we found no evidence for anxiety-like behavior in Open Field or elevated zero maze, and no impairments of short-term memory in Y maze. Further, locomotor behavior was not affected in any of the behavioral assays we performed. These results emphasize the lack of gross behavioral abnormalities in these mice.

Strikingly, however, there were marked impairments of long-term memory in tests of object location memory (**Figure 2A,B**) and contextual fear conditioning (FC, **Figure 2C,D**) in ACSS2 KO mice with AD-tau injection. These tests measure the recall a spatial memory 24 hours after conditioning; OLM requires the mouse to remember an initial arrangement of items in order to identify a displaced object (**Figure 2A**), while in FC, associative memories form between an aversive stimulus (foot shock) and spatial cues from a novel experimental chamber (**Figure 2C**). ACSS2 KO mice^12^ and mice with dorsal hippocampal knock-down of ACSS2^11^ have previously been shown to display impaired performance in these tasks.

With FC, we found that both Tau injection as well as ACSS2 KO resulted in diminished recall of contextual cues (one-way ANOVA, F_3,80_=4.001, p=0.0104; **Figure 2D**). Post-hoc multiple comparisons indicated that the group differences were mostly driven by a significant decrease of freezing in Tau-injected ACSS2 KO mice compared to PBS-injected WT mice (Fisher’s exact, p=0.0049). This finding suggests that the “double hit” of hippocampal Tau injection and loss of ACSS2 significantly worsens long-term memory in these mice.

This pattern of long-term memory deficits was corroborated in a cohort of mice that underwent OLM (**Figure 2A**). The PBS-injected WT mice favored exploration of the object that had been moved on the test day, demonstrating intact long-term spatial memory. However, the discrimination index was markedly decreased in Tau-injected WT and PBS-injected ACSS2 KO mice (**Figure 2B**), indicating long-term memory deficit. As seen in FC, recall was further impacted by the “double hit” of Tau injection and ACSS2 loss (one-way ANOVA F_3,67_=3.281, p=0.0261; Tukey’s test p=0.0310 in the PBS-WT vs AD Tau-KO comparison).

Taken together, these findings suggest that, while individual AD-Tau injection and ACSS2 knock-out both impair long-term memory formation, these deficits are exacerbated with the “double hit” of AD-Tau seeding in animals deficient in ACSS2. As short-term memory, anxiety-like behavior and general locomotion were not affected, these results suggest that loss of ACSS2 selectively contributes to a more severe impairment of long-term memory in this model for AD.

### Epigenetic and transcriptional profiling of AD-Tau injected ACSS2 knock-out mice identifies Rfx3 as a master regulator of gene expression changes

To further define the role of ACSS2 in this model, we next injected AD-Tau into the hippocampus of ACSS2 knock-out (KO) mice^12^ and wild-type (WT) littermates and characterized AD-associated neuropathological and transcriptional changes. Notably, in the ACSS2 KO mouse, AD-Tau seeding induced hyperphosphorylation of endogenous Tau to a similar extent as in WT mice (**Figure 3A**). Intriguingly, however, the combination of AD-Tau injection and loss of ACSS2 led to a striking increase of transcriptional dysregulation when measured by RNA-seq. While there was only a small number of differentially expressed genes (DEGs) either in AD-Tau injection in WT mice or in the ACSS2 KO mice, comparison of WT PBS-injected mice to ACSS2 KO Tau-injected littermates (“double hit” of AD-Tau seeding and ACSS2 loss) revealed over 1,000 DEGs (**Figure 3B**). Gene Ontology analysis showed that dysregulated genes were related to nervous system development, axon guidance, neuronal apoptosis and ion channels (**Figure 3C**), suggesting a broad impairment of neuronal function. In line with findings linking ACSS2 to the regulation of immediate early genes^7,8^, this family of genes was strongly represented among the DEGs, with *Npas4* being the most significantly affected transcript other than *Acss2* itself.

**Figure 3.**
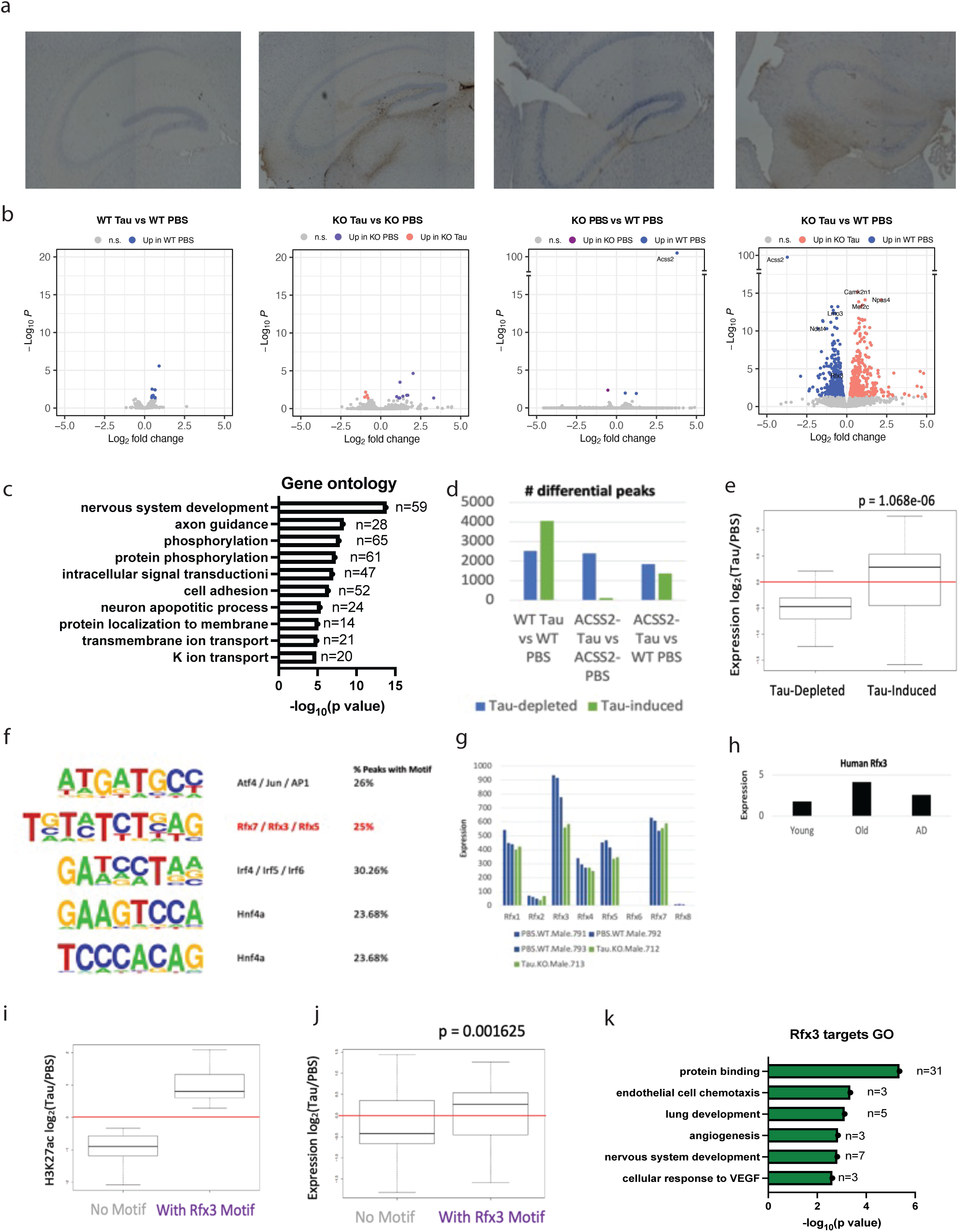
Epigenetic and transcriptional profiling of the AD-Tau injected ACSS2 knock-out hippocampus. **A.** AT8 immunohistochemistry showing similar levels of endogenous Tau phosphorylation in AD-Tau injected WT and ACSS2 KO hippocampi. **B.** Bulk RNA-seq contrasts showing severe transcriptional dysregulation in the hippocampus of AD-Tau injected ACSS2 KO mice. **C.** Gene Ontology analysis of differentially expressed genes in AD-Tau injected ACSS2 KO mice compared to PBS-injected WT controls. **D.** Number of differential H3K27ac peaks in different comparisons. **E.** Expression of genes near H3K27ac peaks induced or depleted by AD-Tau injection. **F.** Motif analysis reveals the presence of Rfx3 binding sites at AD-Tau induced peaks in ACSS2 knock-out hippocampus. **G.** Rfx3 mRNA expression is significantly decreased in the hippocampus of AD-Tau injected ACSS2 knock-out mice, in contrast to other members of the Rfx family. **H.** RFX3 mRNA expression is decreased in the human AD brain compared to healthy age-matched controls. **I.** H3K27ac enrichment is significantly increased at Rfx3 binding sites in AD-Tau injected ACSS2 KO mice. **J.** The expression of genes near Rfx3 binding sites is significantly increased by AD-Tau injection in ACSS2 KO mice. **K.** Gene Ontology analysis of Rfx3 target genes induced by AD-Tau in ACSS2 KO hippocampus.

We then profiled H3K27ac genome-wide by chIP-seq in AD-Tau- or PBS-injected ACSS2 KO mice and WT littermates. There were thousands of differential peaks and Tau-injection increased histone acetylation in WT mice (**Figure 3D left**), which may reflect similar histone acetylation increases at disease-driving genes as we detected in human post-mortem brain^3,4^. In stark contrast, in the ACSS2 KO background, loss of H3K27ac peaks predominated (**Figure 3D middle**). As anticipated, there was a significant association of H3K27ac peaks with gene expression (**Figure 3E**): the expression of genes linked to H3K27 loss was decreased, while expression of genes associated with H3K27 peak gains, increased.

Given the exacerbated behavioral **(Figure 2**) and transcriptional impairments in the WT PBS vs ACSS2 KO AD-Tau injected mice (**Figure 3B**), we focused on this comparison. We investigated potential transcription factors that may be driving the strong gene expression changes with injection of AD-tau into the ACSS2 KO mice. *De novo* motif analysis of differential peaks in these “double hit” mice identified an enrichment of motifs related to AP-1 transcription factor and the family of Rfx transcriptional repressors (**Figure 3F**), with these motifs present in 26% and 25% of all differential peaks, respectively. Remarkably, the expression of *Rfx3*, a gene with frameshift mutations previously linked to neurodevelopmental disorders^35^, was significantly decreased in the hippocampus of the “double hit” mice (**Figure 3G**). *RFX3* expression is also decreased in the post-mortem brain of human AD patients^3,4^ (**Figure 3H**). Strikingly, H3K27ac enrichment (**Figure 3I**) and gene expression (**Figure 3J**) of Rfx3 targets were also significantly increased in the “double hit” mice. Rfx3 target genes are not typically related to neuronal function: top enriched GO categories included protein binding, endothelial cell chemotaxis and lung development (**Figure 3K**). Thus, aberrant de-repression of Rfx3 target genes in AD-Tau-injected ACSS2 KO mice might be a mechanism contributing to impairments of neuronal and cognitive functions (see **Figure 2**) observed in this model.

### Cell-type specific profiling of gene expression and chromatin accessibility highlights impairments of Cajal-Retzius cells in AD-Tau-injected ACSS2 KO mice

Our results highlighted significant behavioral, transcriptional and epigenetic changes in the hippocampi of ACSS2 KO mice in response to AD-Tau-injection, along with potential aberrant transcription factor activity. To determine if there were specific cell populations most affected by AD-Tau and loss of ACSS2, we used the 10xGenomics MultiOmics platform to assess transcription (RNA-seq) and chromatin accessibility (ATAC-seq) from single nuclei. A total of 57,681 nuclei were sequenced from the hippocampi of PBS- or Tau-injected ACSS2 KO mice and WT littermates. Using this strategy, 22 clusters of nuclei were identified that correspond to 10 major neuronal and non-neuronal cell types in the hippocampus (**Figure 4A**). The majority of cell types were relatively stable across the four conditions; however, we observed a diminished number of pyramidal neurons and Cajal-Retzius (CR) neurons in AD-Tau-injected mice, and reduced astrocyte numbers in AD-Tau-injected ACSS2 KO mice (**Figure 4B**). The CR cells were especially intriguing as they showed the largest transcriptional impairments when we comparing WT mice to AD-Tau-injected ACSS2 KO animals (**Figure 4C**). CR cells are glutamatergic neurons in the cortex and hippocampus that play an important role developmentally and have been linked to learning and memory in adult animals^36,37^. Loss of CR cells at different developmental stages has been reported in transgenic mouse models of AD^38,39^. Here, 1,256 differentially expressed genes were identified in CR neurons in the WT PBS *vs*. KO AD-Tau comparison. Out of all clusters and cell types, the DEGs observed in CR cells showed the most significant overlap and strongest positive correlation with DEGs from bulk RNA-seq (R = 0.4, **Figure 4D,E**) emphasizing the important overall influence of transcriptional dysregulation in this cell type on the observed hippocampal gene expression changes. As with the bulk RNA-seq (**Figure 3**), transcriptional changes were present but less pronounced with AD-Tau injection only (361 DEGs) or ACSS2 KO only (458 DEGs). Remarkably, the pattern of gene expression changes in CR cells matched the severity of behavioral impairments in the fear conditioning assay (see **Figure 2**): the top upregulated and top downregulated genes were increased or decreased both by AD-Tau injection and loss of ACSS2, with significantly more pronounced changes in the “double hit” mice (**Figure 4F**). GO analysis indicated enrichment of genes related to synaptic functions including synapse organization and regulation of membrane potential (**Figure 4G**), suggesting that the activity of CR cells is severely compromised in AD-Tau-injected ACSS2 KO mice. Moreover, several of the top DEGs have been linked to AD or other forms of neurodegeneration. For example, the transcription factor *Tshz2* (Teashirt zinc finger homeobox 2) is increased in excitatory neurons of AD patients^40^, while ETS variant transcription factors such as *Etv1* are linked to AD pathogenesis via elevated BACE1 levels, which increases cleavage of the amyloid precursor protein^41^.

**Figure 4.**
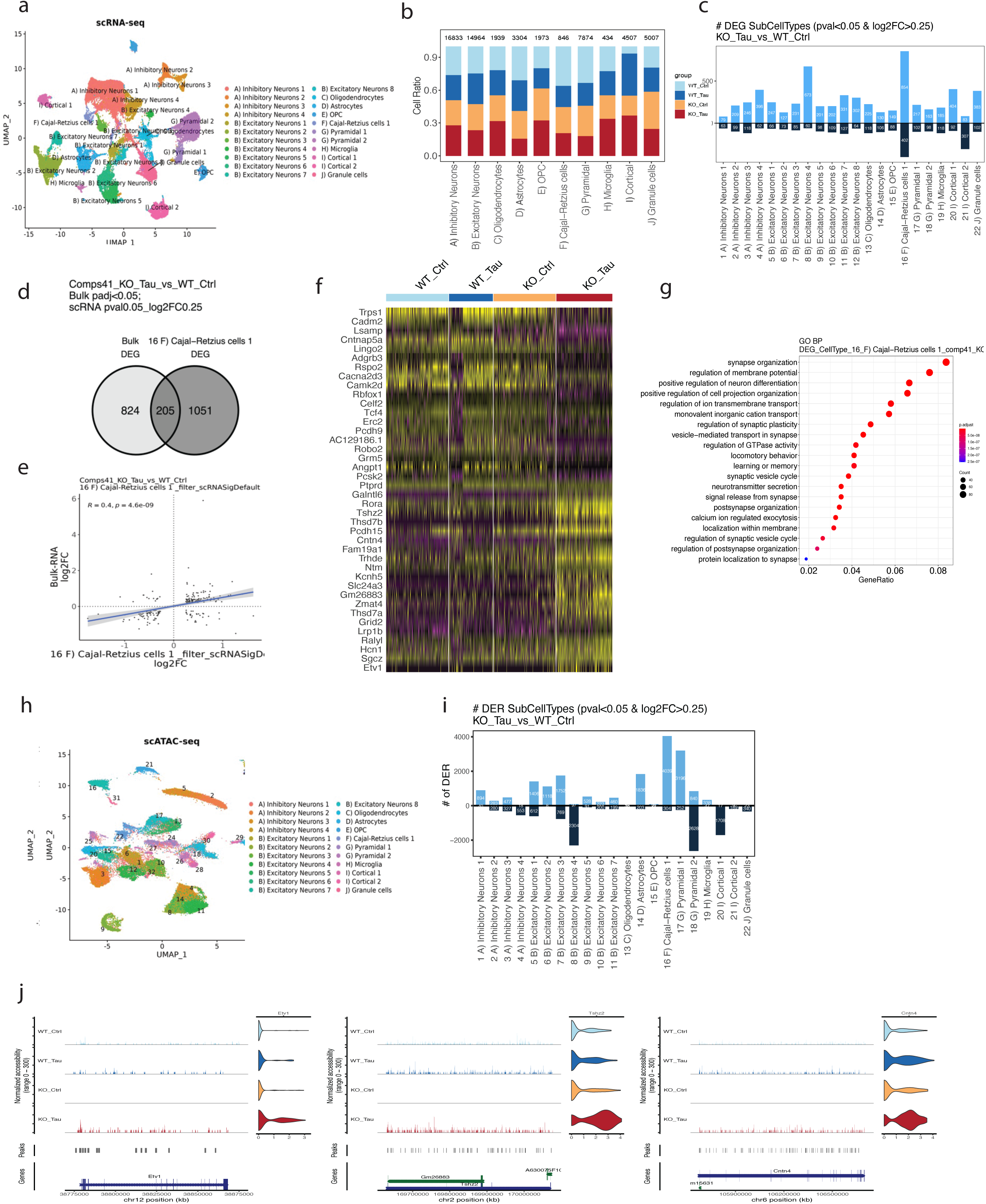
Single nuclei transcriptional and chromatin profiling highlights impact on Cajal-Retzius cells. **A.** UMAP embedding of neuronal and non-neuronal cell types identified in single nuclei RNA-seq of mouse hippocampus. **B.** Changes in cell number across conditions (genotype, injection). AD-Tau injection leads to loss of pyramidal cells, astrocytes and Cajal-Retzius (CR) neurons. **C.** CR cells show the largest number of differentially expressed genes across all clusters in AD-Tau injected ACSS2 KO mice compared to PBS-injected WT controls. **D.** Differential genes called in CR cells overlap significantly with differential genes from hippocampal bulk RNA-seq. **E.** Differential genes called in CR cells show a positive correlation with differential genes from hippocampal bulk RNA-seq. **F.** Heatmap of differential genes in CR cells across conditions reveals more severe impairment of gene expression in AD-Tau injected ACSS2 KO mice compared to other groups. **G.** Gene Ontology analysis of differentially expressed genes in CR cells of AD-Tau injected ACSS2 KO mice (circle size = # genes; heat = significance p-value). **H.** UMAP embedding of neuronal and non-neuronal cell types identified in single nuclei ATAC-seq of mouse hippocampus. **I.** CR cells show the largest number of differentially accessible regions across all clusters in AD-Tau injected ACSS2 KO mice compared to PBS-injected WT controls. **J.** Chromatin accessibility at top differentially expressed genes in CR cells.

The impairment of CR cells was supported by chromatin landscape changes defined by snATAC-seq. We examined DNA from 47,672 nuclei to measure chromatin accessibility, and used markers from the snRNA-seq data to annotate clusters (**Figure 4H**). Overall, the clusters identified by snRNA-seq and snATAC-seq integrated well and chromatin accessibility was a good predictor of gene expression in different cell types. CR cells again exhibited the largest number (4,343) of differentially accessible regions when we compared WT PBS to KO AD-Tau injected mice (**Figure 4I**). Of note, this included increased chromatin accessibility of top upregulated genes, including *Tshz2* and *Etv1* (**Figure 4J**). Taken together, the multiomic profiling provided converging evidence for a robust impact on Cajal-Retzius neurons, emphasizing a potential key role of this hippocampal cell type in Alzheimer’s disease.

### Acetate-enriched diet ameliorates AD-Tau induced molecular changes and memory deficits

Since inhibiting metabolic-epigenetic interactions by knocking out the nuclear metabolic enzyme ACSS2 exacerbates molecular and behavioral phenotypes observed in the mouse AD-tau model, we considered that stimulating this pathway by supplying the substrate of ACSS2, acetate, might have beneficial effects. Acetate supplementation to primary hippocampal neurons drives genes of learning and memory and promotes cognitive function in a developmental mouse model^42^, and acetate gavage can promote memory in the 5xFAD mouse^43^. To investigate and define mechanisms of acetate supplementation in modulation of gene function for learning and memory in normal mice, first we tested whether exogenous acetate is directly incorporated into hippocampal histone acetylation, using heavy labeled acetate. Mice were injected with d3-acetate i.p., and heavy labeling of acetylated histones was measured at different timepoints using stable isotope labeling mass spectrometry. We observed a transient time-dependent (peak labeling at 30 min) and dose-dependent deposition of acetate on hippocampal histones (**Figure 5A,B**). Heavy label incorporation was nearly completely dependent on ACSS2; heavy label was not incorporated into HPC histones in ACSS2 KO mice (**Figure 5C**).

**Figure 5.**
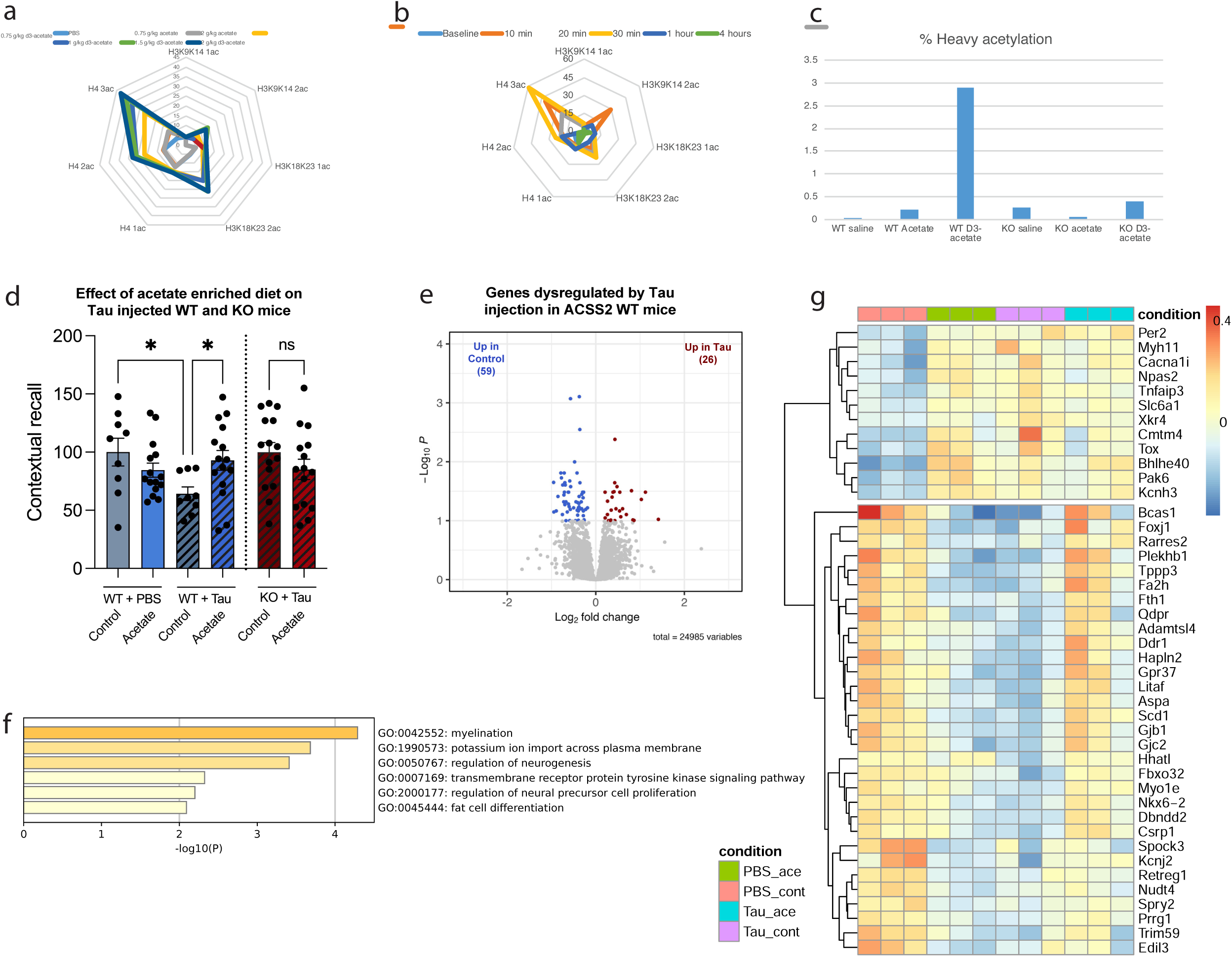
Acetate-enriched diet ameliorates AD-Tau induced molecular changes and memory deficits. **A.** Dose-dependent heavy labeling of mouse hippocampal histone acetylation following d3-acetate injection. **B.** Temporal dynamics of heavy labeling of mouse hippocampal histone acetylation following d3-acetate injection. **C.** Heavy labeling of histone acetylation in WT but not ACSS2 KO mice following d3-acetate injection. **D.** Dietary supplementation of acetate increases freezing behavior during recall of contextual fear memories in WT mice, but not ACSS2 KO mice. **E.** Volcano plot showing genes differentially regulated in Tau-injected WT mice compared to PBS-controls. **F.** Gene Ontology analysis of genes downregulated in Tau-injected WT mice. **G.** Heatmap of top 45 differentially regulated genes in Tau-injected WT mice compared to PBS-controls, showing partial rescue of downregulated transcripts with acetate supplementation.

We then investigated whether stimulating ACSS2-driven histone acetylation via supplying exogenous acetate can reverse learning and memory impairments in the context of AD using AD-tau injection. WT mice were injected with 1 μg AD-Tau in the dorsal hippocampus and underwent fear conditioning 6 months post-injection. Acute exposure to acetate (1.5 g/kg i.p. prior to acquisition) did not rescue decreased freezing in this model, thus we hypothesized that chronic administration of acetate might be required to reverse the long-lasting effects of AD-Tau seeding. To achieve chronic acetate administration, we maintained mice on a special diet enriched in acetate (5% w/w) throughout the 6 months period post Tau-injection. As previously reported^44^, mice administered the acetate diet had slightly reduced body weight compared to controls maintained on a calorically equivalent diet (Student’s t test, t_46_=2.236, p=0.0302), but we found no signs of sickness, anxiety or any impairments of general locomotion in Open Field. Chronic acetate supplementation also had no effect on freezing behavior in mice that were not AD-Tau-injected (Student’s t test, t_22_=1.276, p=0.2153). In AD-Tau-injected mice, however, the acetate diet dramatically improved contextual recall, evidenced by a nearly 50% increase in freezing behavior (Student’s t test, t_22_=2.397, p=0.0254) (**Figure 5D, left**). Notably, this ameliorative effect was only observed in WT mice: acetate-enriched diet did not improve recall of fear-associated contextual cues in ACSS2 KO mice (Student’s t test, t_28_=1.238, p=0.2261) (**Figure 5D, right**).

Next, we examined the transcriptional landscape underlying the acetate-mediated rescue of Tau-related deficits. WT mice were seeded with 1 μg AD-Tau and maintained for 6 months on an acetate-rich or control diet. Control mice were sham injected with PBS. We performed bulk RNA-seq on the dorsal hippocampus of representative animals from each group (n=3), selecting the side where AD-Tau had been seeded 6 months prior. Consistent with previous results (**Figure 3B**), we found that Tau injection led to a modest number of differentially expressed genes in mice fed standard chow (**Figure 5E**). We noted the affected GO terms included neurogenesis and myelination (**Figure 5F**), impairments of which are canonical features of human AD. Strikingly, of the 56 genes downregulated by Tau injection, all showed recovery with acetate supplementation, providing evidence that acetate supplementation can not only enhance long-term memory but also facilitate neuronal resilience and recovery (**Figure 5G**).

Taken together, these data indicate that boosting ACSS2-dependent histone acetylation by dietary supplementation of acetate rescues long-term memory in a mouse model of AD, while having no discernible side effects that would confound its translational potential.

## Discussion

We establish ACSS2-mediated histone acetylation as a novel modulator of Alzheimer’s disease (AD)-related molecular and behavioral phenotypes in an AD-tau seeding mouse model that mimics the progress of human pathology. We report that loss of ACSS2 exacerbates memory impairments in this model (**Figure 2**), while dietary supplementation of acetate, the substrate of this enzyme, rescues learning (**Figure 5**). Further, we show that this effect is mediated by the ACSS2-dependent incorporation of acetate directly into hippocampal histone acetylation, which facilitates gene expression programs that drive learning (**Figure 5**). Characterization of the underlying histone acetylation and transcriptional changes using single nuclei approaches identified Cajal-Retzius neurons as a critical hippocampal neuronal population affected by AD (**Figure 4**). Intriguingly, the epigenetic and gene expression changes manifest separately from hippocampal AD-Tau pathology (**Figure 1**), thus our work outlines new mechanisms underlying the offset and progression of AD.

Tau seeding has been primarily used to study mechanisms that govern the assembly of Tau into filamentous inclusions, as well as the spreading of Tau pathology across brain regions and differences between distinct conformers^19,27^. Notably, here we show that the molecular responses to AD-Tau injection reflect the molecular characteristics of human AD (**Figure 3**). Specifically, histone acetylation and transcriptional changes induced by AD-Tau injection mirror those previously reported in the brain of AD patients^3,4^. Genes associated with the top AD-Tau-induced H3K27ac peaks showed a striking enrichment in genes dysregulated in human AD^28,29^; several differentially expressed genes are linked to AD risk, neuronal death, and memory loss^31^. Overall, these findings corroborate and underscore AD-Tau injection as a translational mouse model of AD that, in addition to Tau pathology, also recapitulates the epigenetic and transcriptional changes reflective of the brain of human AD patients.

Therapeutic interventions focusing on well-established pathological aspects of AD, such as amyloid plaque formation, continue to show limited success in clinical settings^45^. Ongoing efforts to uncover the molecular underpinnings of AD early in the disease are thus of critical importance. Recently, epigenetic mechanisms, and in particular histone acetylation, have emerged as central players in the development of AD^46^. We and others have reported marked changes of histone acetylation in the human AD brain. For example, acetylation of histone H4 lysine 16 is decreased in human AD brains compared to age-matched controls, while acetylation of histone H3 lysine 9 and of histone H3 lysine 27 is increased^3,4^. This intriguing dichotomy of protective and risk-conferring histone acetylation is potentially related to distinct biological functions associated with different lysine residues^13^. H3K9ac and H3K27ac are activating marks, and their increased enrichment drives gene expression programs related to disease pathways and the dysregulation of transcription and chromatin feedback loops. On the other hand, H4K16ac is a reservoir mark, which has been shown to increase during normal aging in humans and in various model organisms: the transfer of acetyl groups from reservoir to activating sites is a novel form of epigenetic regulation^13^ that might be instrumental for rapid transcriptional adaptations required for neuronal activity. The loss of H4K16ac and other reservoir marks in AD could result in decreased transcriptional plasticity of neurons, which is a critical molecular mechanism required for learning and memory.

Despite the emergence of histone acetylation as a key epigenetic mechanism contributing to AD, therapeutic approaches targeting this pathway in AD have been challenging. Inhibitors of histone deacetylases (HDACs) and histone acetyltransferases (HATs) have off-target effects, and the HDAC and HAT enzymes themselves can be non-specific. Similarly, the therapeutic use of bromodomain inhibitors is precluded by the toxic side effects of these drugs. Recent evidence is pointing to the role of nuclear metabolic enzymes as key regulators of epigenetic mechanisms prominently including histone acetylation. These enzymes are directly recruited to the nucleus and chromatin, where they generate a local, transient supply of metabolites as substrates or cofactors for histone and DNA modifiers^29^. ACSS2, for example, utilizes acetate to generate acetyl-CoA which is deposited by histone acetyltransferases, thereby increasing the expression of genes related to neuronal activity and memory formation^7^. This metabolic-epigenetic interface provides an intriguing avenue for therapeutic interventions. Upregulation of ACSS2 and acetate gavage have been shown to improve learning and memory for the Morris water maze in an APP mouse model^43^. Strikingly, here we show that simple dietary supplementation of acetate rescues memory formation in the AD-tau injection mouse—an effect that is dependent on ACSS2-mediated histone acetylation in the hippocampus. Consistent with a key role for ACSS2, inhibition of ACSS2 by genetic deletion severely exacerbates learning deficits in mice injected with AD-Tau. Importantly, these manipulations had no effect on short-term memory, anxiety-like behavior or general locomotion, and thus hold promise for increased specificity and lower toxicity in a clinical setting. Further, dietary supplementation of acetate is entirely non-invasive and would likely pose limited compliance issues during chronic administration.

Our single nuclei RNA-seq and ATAC-seq data highlight the surprising importance of Cajal-Retzius cells (CR), a small subpopulation of hippocampal neurons, in AD. CR cells are glutamatergic neurons in the cortex and hippocampus that are primarily studied in the context of brain development, although have also been linked to learning and memory^36,37^. Loss of CR cells at various developmental time points has been observed in a transgenic mouse model of AD^38,39^. Here we find that CR cells show the largest number of differentially expressed genes and differentially accessible chromatin regions, in addition to a decrease in their cell number following AD-Tau injection and ACSS2 knock out. Despite the relatively small population of CR cells in the adult mouse hippocampus, our findings highlight that this cell type has a striking influence on the epigenetic and transcriptional landscape of this brain region overall, as evidenced by the significant overlap and positive correlation between differential genes called from CR cells and bulk hippocampal RNA-seq. Intriguingly, the number of CR cells shows remarkable plasticity and increases upon environmental enrichment^47^, which has beneficial effects on cognition. Whether dietary supplementation of acetate exerts its therapeutic effects by acting on CR cells will be an important area for future investigation.

Taken together, we establish AD-Tau injection as a mouse model that reflects molecular changes observed in human AD. We show that epigenetic and transcriptional changes are exacerbated following ACSS2 knock out and are independent from Tau pathology, and we identify Cajal-Retzius cells as key contributors to the molecular re-configuration of the AD hippocampus. Further, we show that genetic deletion of ACSS2 significantly worsens long-term memory in the context of Tau injection, pointing to an important neuroprotective role of ACSS2-mediated histone acetylation in the hippocampus. Indeed, dietary supplementation of acetate, the substrate of ACSS2, rescued learning and memory in an ACSS2-dependent manner. Overall, this work establishes ACSS2 and metabolic-epigenetic dysregulation as a novel aspect of AD-associated neurobiology, with the potential to inform the development of future therapies.

